## Supplemental Figures for "Genome-wide Bioinformatics Analysis of Human Protease Specificity Identified Potential Cathepsin L Cleavage Site at K790 Position of the SARS-CoV-2 Spike Glycoprotein"

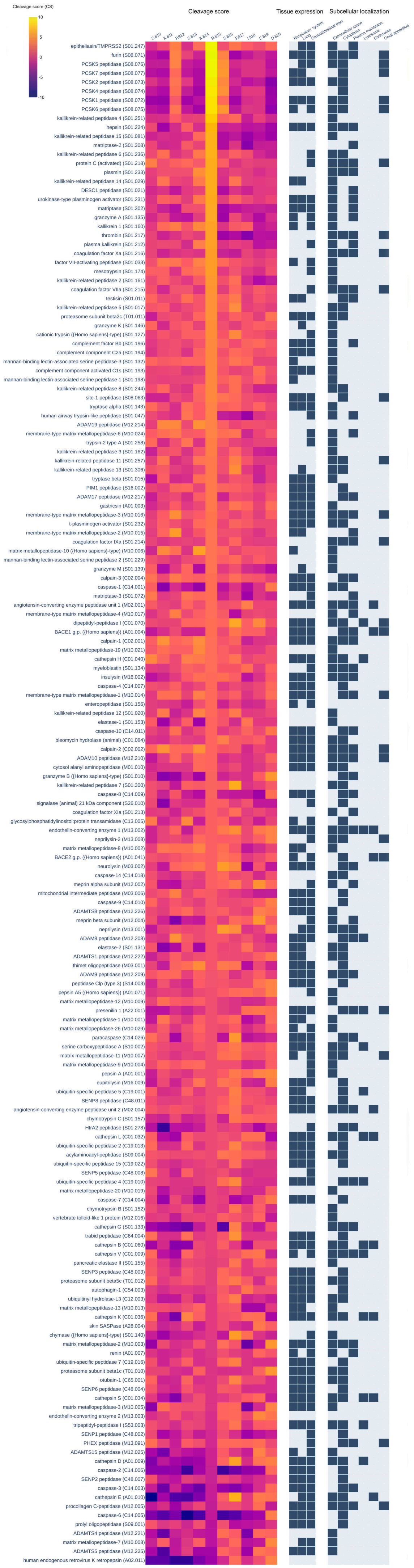


**Figure S1.** Cleavage scores at the R815 position, protease cellular localization, and protease tissue expression for the 169 proteases with modeled protease sequence specificity.

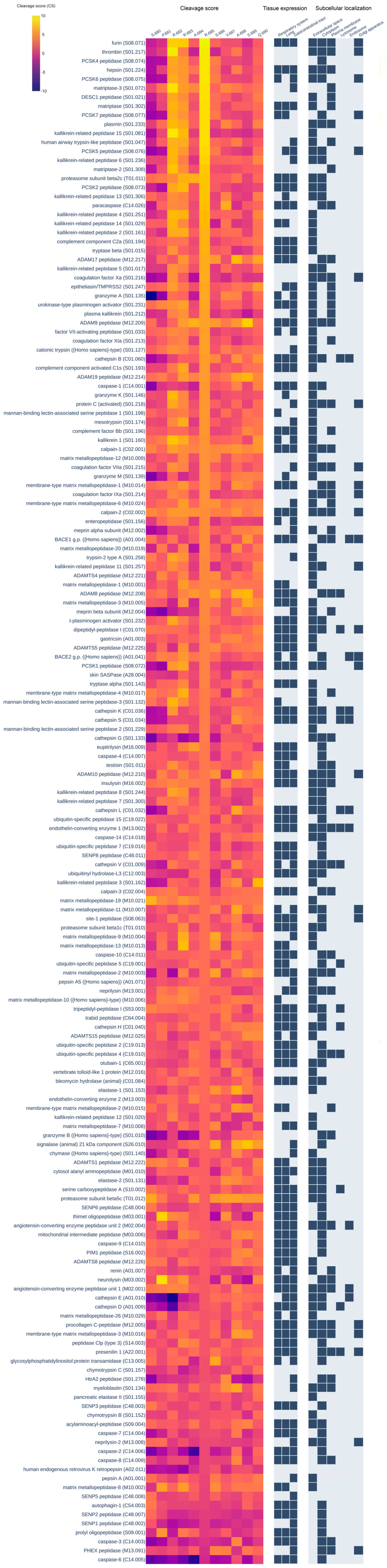


**Figure S2.** Cleavage scores at the R685 position, protease cellular localization, and protease tissue expression for the 169 proteases with modeled protease sequence specificity.

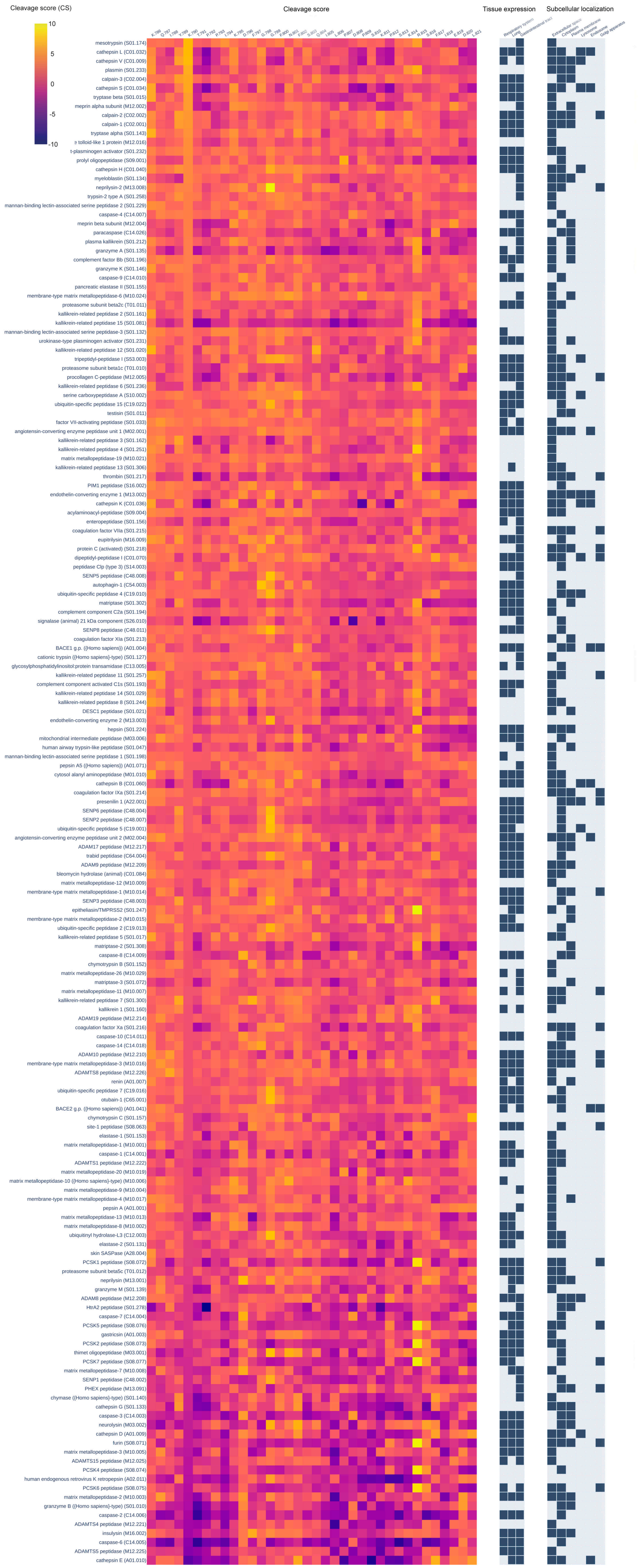


**Figure S3.** Cleavage scores at the K790 position, protease cellular localization, and protease tissue expression for the 169 proteases with modeled protease sequence specificity.

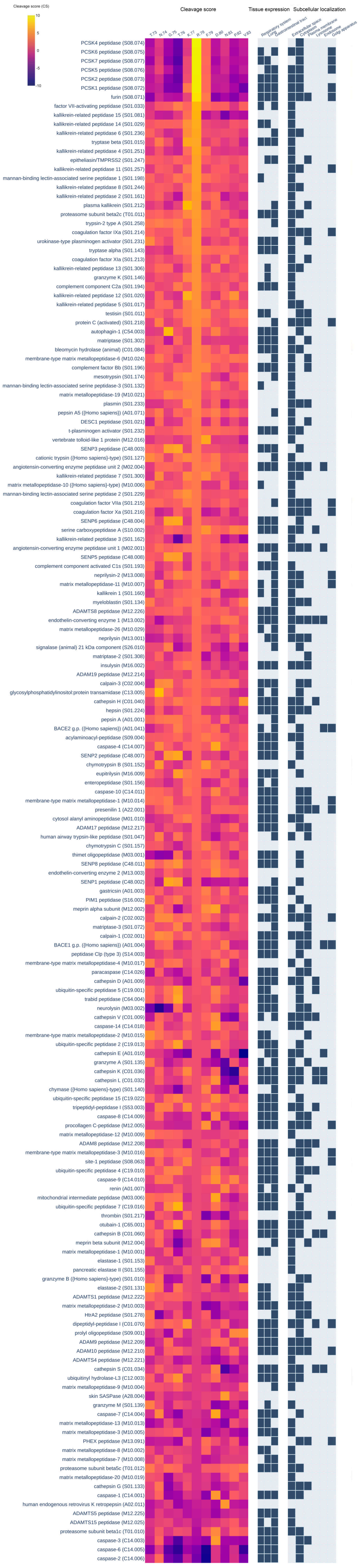


**Figure S4.** Cleavage scores at the R78 position, protease cellular localization, and protease tissue expression for the 169 proteases with modeled protease sequence specificity.

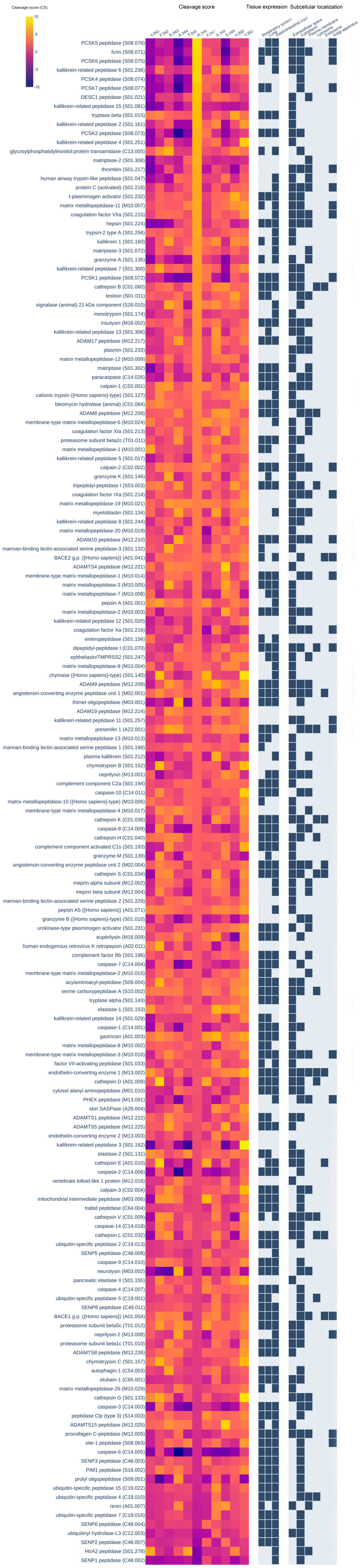


**Figure S5.** Cleavage scores at the R346 position, protease cellular localization, and protease tissue expression for the 169 proteases with modeled protease sequence specificity.

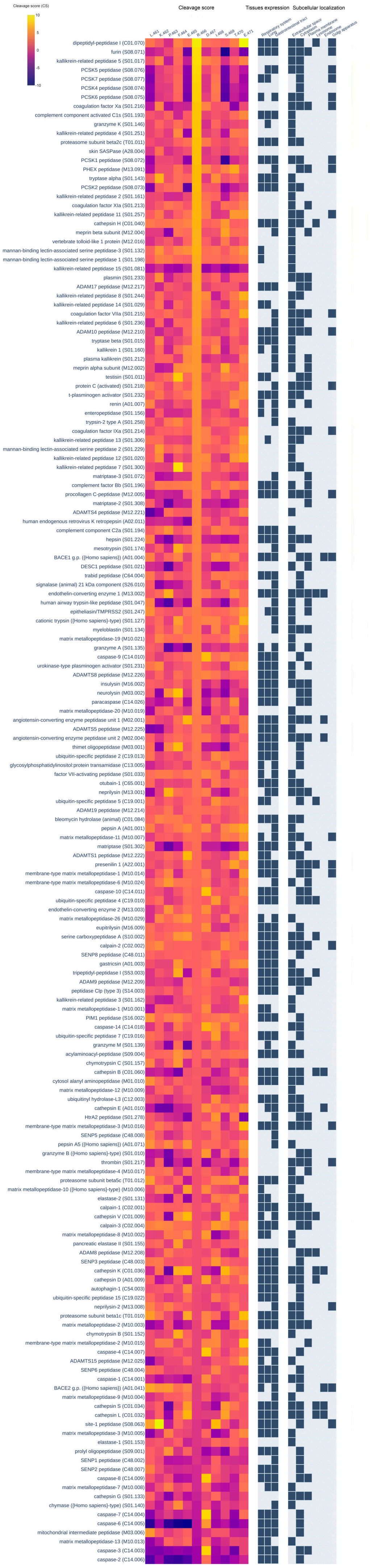


**Figure S6.** Cleavage scores at the R466 position, protease cellular localization, and protease tissue expression for the 169 proteases with modeled protease sequence specificity.

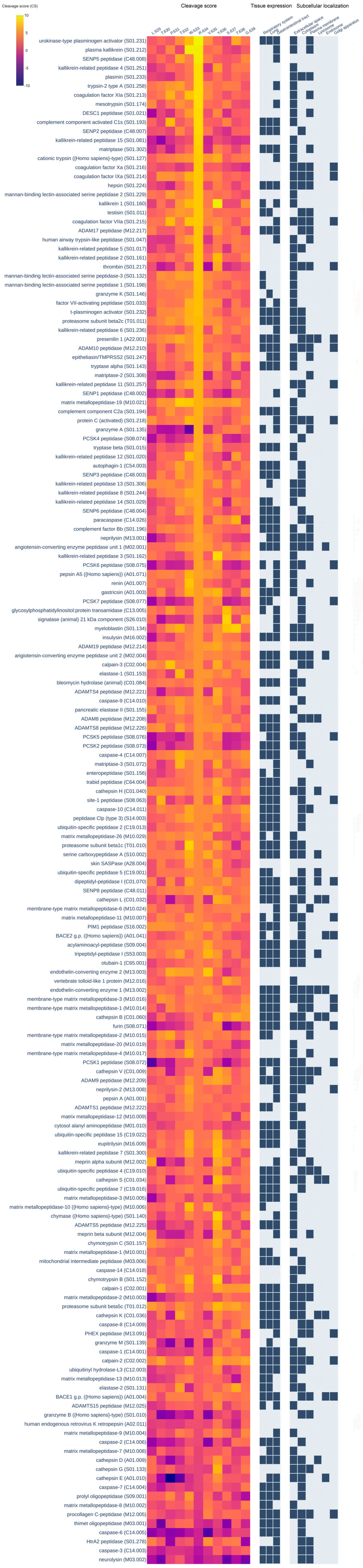


**Figure S7.** Cleavage scores at the R634 position, protease cellular localization, and protease tissue expression for the 169 proteases with modeled protease sequence specificity.

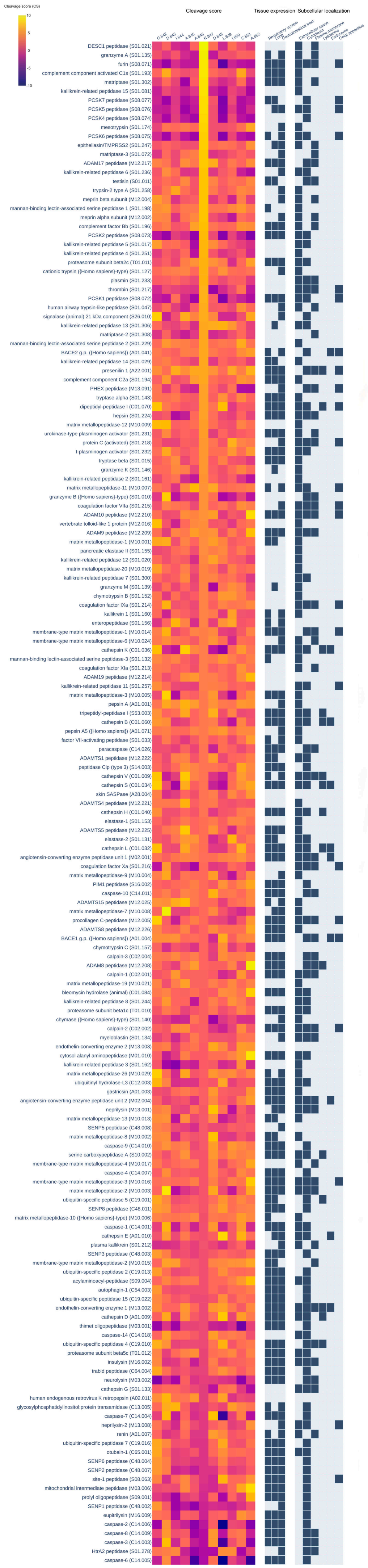


**Figure S8.** Cleavage scores at the R847 position, protease cellular localization, and protease tissue expression for the 169 proteases with modeled protease sequence specificity.

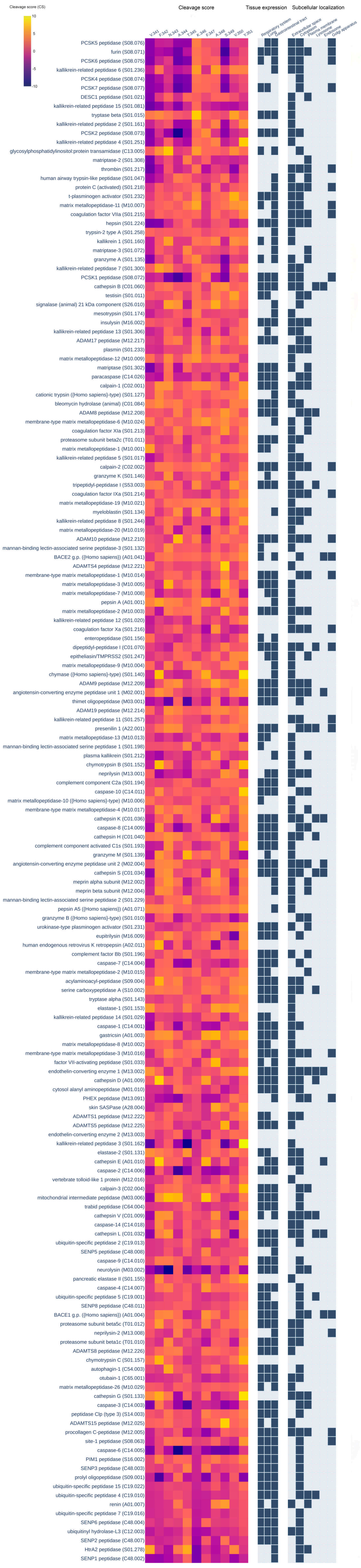


**Figure S9.** Cleavage scores at the R346K position, protease cellular localization, and protease tissue expression for the 169 proteases with modeled protease sequence specificity.

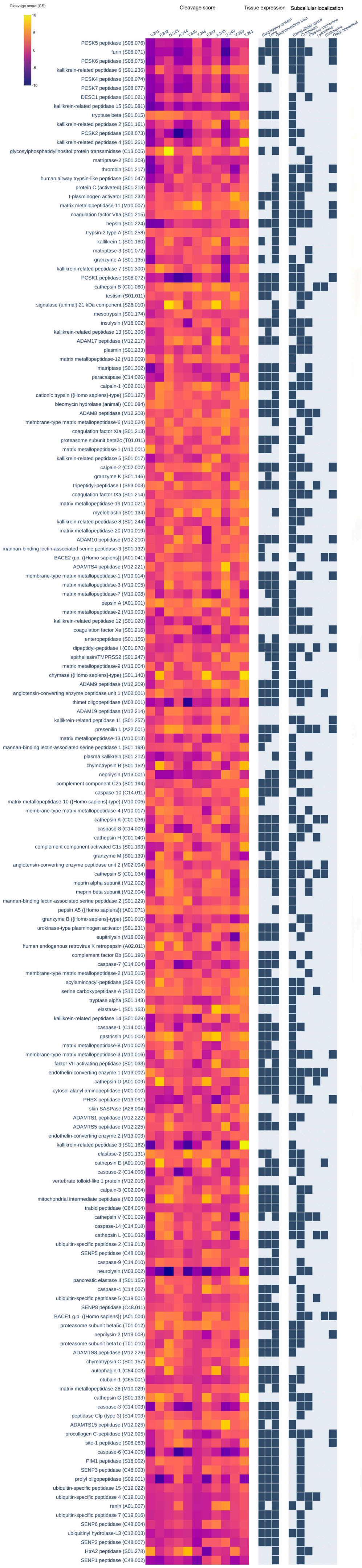


**Figure S10.** Cleavage scores at the R346T position, protease cellular localization, and protease tissue expression for the 169 proteases with modeled protease sequence specificity.

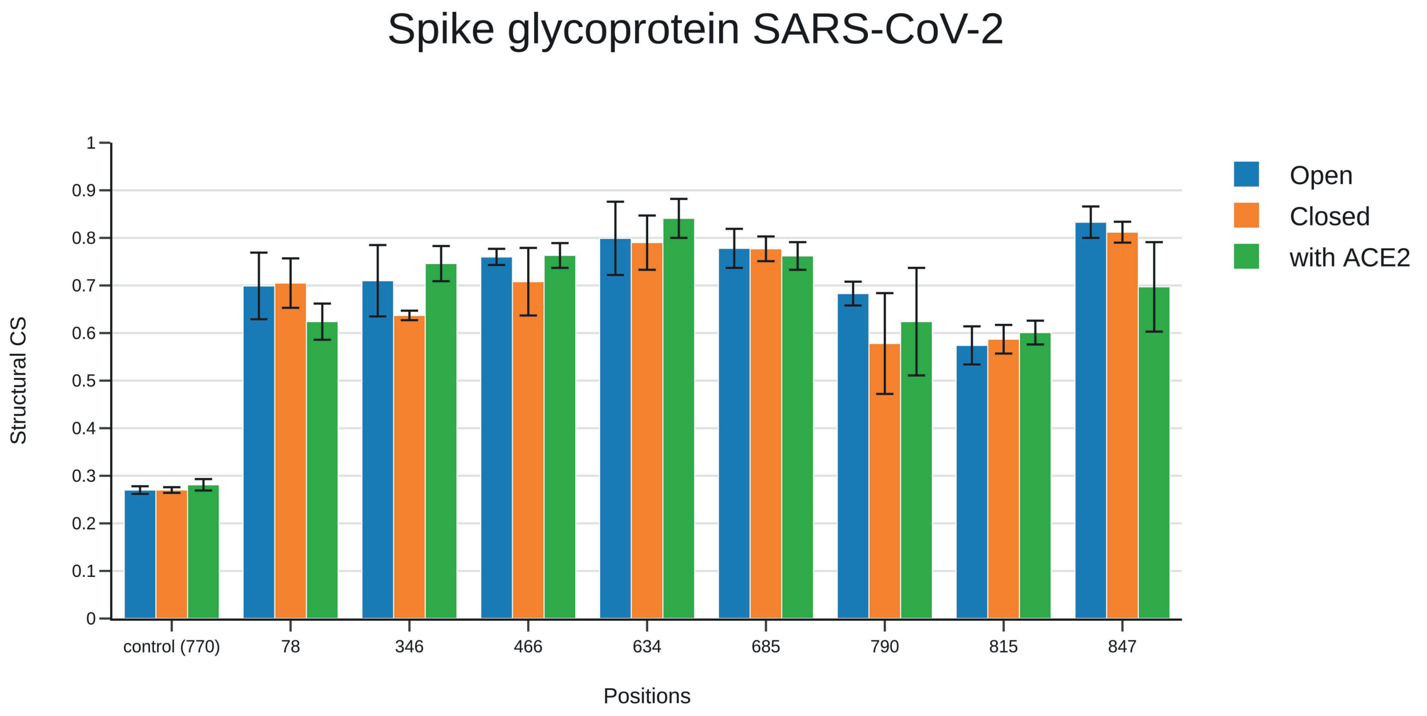


Figure S11. Structural estimation of susceptibility to proteolysis for known and potential cleavage sites of the spike glycoprotein based on 3D structures from different SARS-CoV-2 variants (Alpha, Beta, Delta, Kappa, Omicron).
